# Mendelian randomization analyses support causal relationships between blood metabolites and the gut microbiome

**DOI:** 10.1101/2020.06.30.181438

**Authors:** Xiaomin Liu, Xin Tong, Yuanqiang Zou, Xiaoqian Lin, Hui Zhao, Liu Tian, Zhuye Jie, Qi Wang, Zhe Zhang, Haorong Lu, Liang Xiao, Xuemei Qiu, Jin Zi, Rong Wang, Xun Xu, Huanming Yang, Jian Wang, Yang Zong, Weibin Liu, Yong Hou, Shida Zhu, Huijue Jia, Tao Zhang

**Affiliations:** BGI-Shenzhen, Shenzhen, China; College of Life Sciences, University of Chinese Academy of Sciences, Beijing 100049, China; China National Genebank, BGI-Shenzhen, Shenzhen, China; Shenzhen Engineering Laboratory of Detection and Intervention of Human Intestinal Microbiome, BGI-Shenzhen, Shenzhen, China; BGI-Qingdao, BGI-Shenzhen, Qingdao, China; James D. Watson Institute of Genome Sciences, Hangzhou, China; Shenzhen Key Laboratory of Human Commensal Microorganisms and Health Research, BGI-Shenzhen, Shenzhen, China

## Abstract

The gut microbiome has been implicated in a variety of physiological states, but controversy over causality remains unresolved. Here, we performed bidirectional Mendelian randomization (MR) analyses on 3,432 Chinese individuals with whole genome, whole metagenome, anthropometric, and blood metabolic trait data. We identified 58 causal relationships between the gut microbiome and blood metabolites, and replicated 43 of them. Increased relative abundances of fecal *Oscillibacter* and *Alistipes* were causally linked to decreased triglyceride concentration. Conversely, blood metabolites such as glutamic acid appeared to decrease fecal *Oxalobacter*, and members of *Proteobacteria* were influenced by metabolites such as 5-methyltetrahydrofolic acid, alanine, glutamate, and selenium. Two-sample MR with data from Biobank Japan partly corroborated results with triglyceride and with uric acid, and also provided causal support for published fecal bacterial markers for cancer and cardiovascular diseases. This study illustrates the value of human genetic information to help prioritize gut microbial features for mechanistic and clinical studies.

---

Metagenome-wide association studies (MWAS) using human stool samples, as well as animal models, especially germ-free mice, have pointed to a potential role of the gut microbiome in diseases such as cardiometabolic, autoimmune, neuropsychiatric disorders and cancer, with mechanistic investigations for diseases such as obesity, colorectal cancer and schizophrenia^1–4^. Twin-based heritability estimation and more recent metagenome-genome-wide association studies (M-GWAS) have questioned the traditional view of the gut microbiota as a purely environmental factor^5–9^, although the extent of the genetic influence remains controversial^7,10^. Yet, all these published cohorts, except for human sequences in the metagenomic data of HMP (Human Microbiome Project), utilized array data for human genetics, and most of them had 16S rRNA gene amplicon sequencing for the fecal microbiota^5–9^.

As the gut microbiome is considered to be highly dynamic, causality has been an unresolved issue in the field. Mendelian randomization (MR)^11^ offers an opportunity to distinguish between causal and non-causal effects from cross-sectional data, without animal studies or randomized controlled trials. An early study used MR to look at the gut microbiota and ischemic heart disease^12^. Recently, a study used MR to confirm that increased relative abundance of bacteria producing the fecal volatile short-chain fatty acid (SCFA) butyrate was causally linked to improved insulin response to oral glucose challenge; in contrast, another fecal SCFA, propionate, was causally related to an increased risk of T2D^13^. However, both studies used genotype data, and it was not clear to what extent the genetic factors explained the microbial feature of interest.

In this study, we present a large-scale M-GWAS using whole genome and fecal microbiome, followed by bidirectional MR for the fecal microbiome and anthropometric features as well as blood metabolites. In a two-stage design from different cities in China, we identified 58 causal links from MR in the 4D-SZ discovery cohort of 2,002 individuals with high-depth whole-genome sequencing data (1,539 individuals with microbiome data for one-sample MR). We replicated 43 of the 58 causal effects using low-depth whole-genome sequencing data of another 1,430 individuals (1,006 individuals with microbiome data for one-sample MR). In general, unidirectional causal effects could be found both from the gut to the blood and from the blood to the gut, but bidirectional effects were rarely detected. A few of the M-GWAS associations with gut microbial functional modules, e.g. module for lactose/galactose degradation and the *ABO* loci, reached study-wide significance, illustrating the power of shotgun metagenomic data together with whole genome sequencing. The MR findings were corroborated and extended by summary statistics from the Biobank Japan study, e.g. causal effect of *Proteobacteria* on type 2 diabetes (T2D), congestive heart disease and colorectal cancer, underscoring the significance of human genetic data to help guide microbiome intervention studies.

## Results

### Fecal microbiome features associated with human genetic variants

We set out to identify human genetic variants to be included as the randomizing layer of MR (**Fig. 1**). The 4D-SZ (multi-omics, with more time points to come, from Shenzhen, China) discovery cohort consisted of high-depth whole-genome sequencing data from 2,002 blood samples (mean depth of 42×; **Supplementary Table 1** and **Supplementary Fig. 1a**), out of which 1,539 individuals had metagenomic shotgun sequencing data from stool samples (8.56 ± 2.28 Gb; **Supplementary Fig. 1b**). Fecal M-GWAS was performed using 10 million common and low-frequency variants (minor allele frequency (MAF) ≥ 0.5%) and 500 unique microbial features (Spearman’s correlation < 0.99; **Methods**). The M-GWAS was adjusted for age, gender, BMI, defecation frequency, stool form, self-reported diet, lifestyle factors, and the first four principal components from the genomic data to account for population stratification.

**Figure 1.**
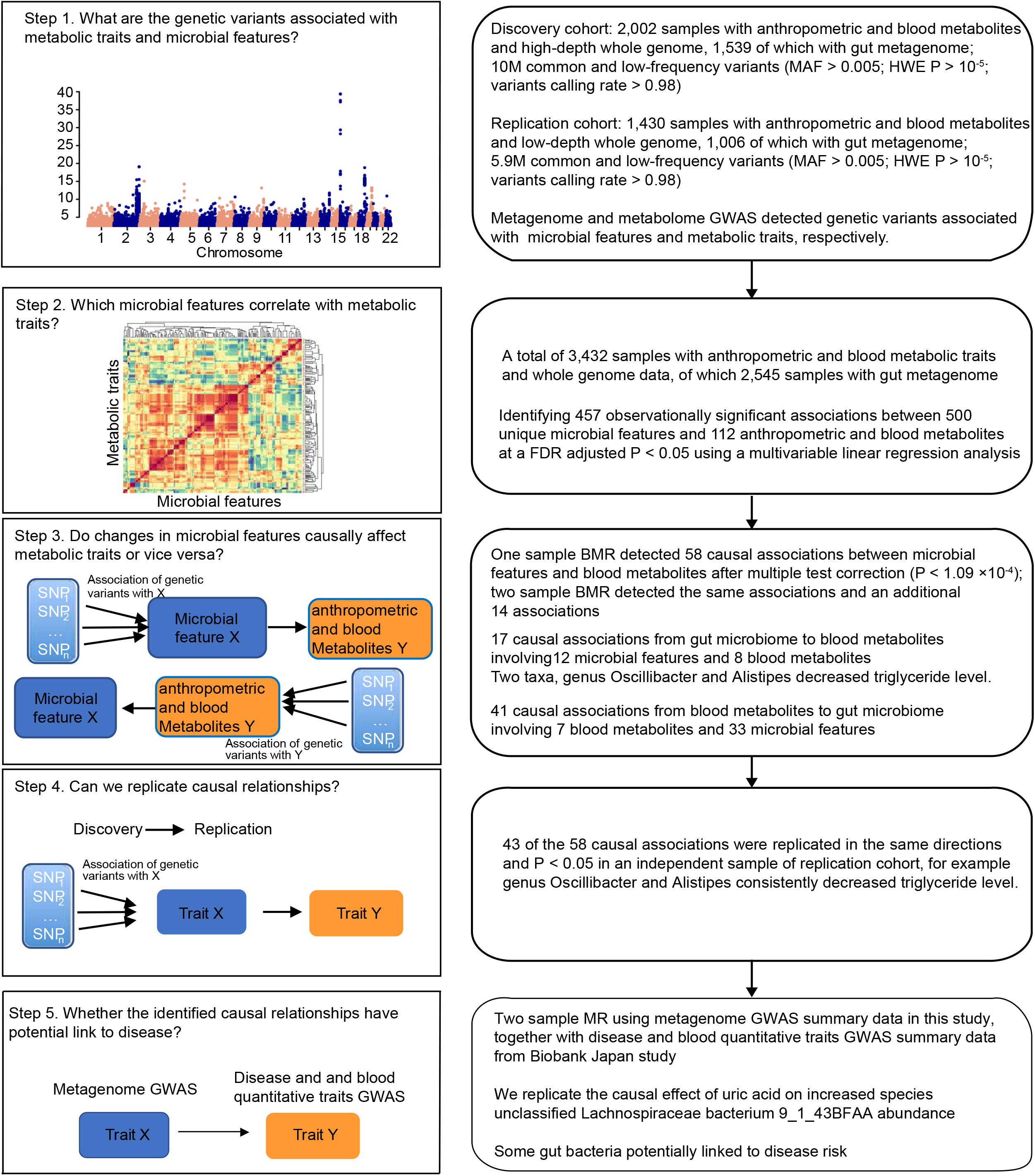
Study design and workflow. This schematic representation highlights, for each step, the research question that we sought to answer, the analysis workflow, the data used, and the generalized result. We first performed metagenome and metabolome GWAS to detect genetic variants associated with microbial features and metabolic traits, respectively, both in discovery and replication cohorts (Step 1). We then performed observational analysis to identify which microbial feature (taxa, GMM) correlated with metabolic traits in this cohort (Step 2). We used 2,545 samples with information of both microbial features and metabolic traits. We observed 457 significant associations between 500 unique microbial features and 112 anthropometric and blood metabolic traits at an FDR adjusted *P* < 0.05. We then estimated causal relationships for the 457 observational associations through bidirectional MR analysis in discovery cohort (Step 3). One-sample BMR detected 58 causal associations between microbial features and blood metabolites after multiple test correction (*P* < 1.09 × 10^−4^); two-sample BMR detected the same associations and an additional 14 associations. As a validation, we replicated the discovered causal relationships by using the same MR analysis in an independent replication cohort (Step 4). Over half (43) of the 58 causal associations were replicated in the same direction (*P* < 0.05). Finally, we used two-sample MR analysis to investigate the effects of the identified 72 causal relationships on diseases from the Biobank Japan study (Step 5).

With this large cohort of whole genome and whole metagenome data, we performed M-GWAS analysis and identified a total of 625 associations involving 548 independent loci for one or more of the 500 microbial features at genome-wide significance (*P* < 5 × 10^−8^). With a more conservative Bonferroni-corrected study-wide significant *P* value of 1.0 × 10^−10^ (= 5 × 10^−8^/500), we identified 28 associations with fecal microbial features involving 27 genomic loci, of which 5 correlated with gut bacteria and the other 22 associated with gut metabolic pathways (**Supplementary Table 2**).

For MR, it was important for the genetic variants used to be representative of the microbiome features (**Supplementary Fig. 2**), so a more suggestive *P* value of less than 1 × 10^−5^ was used (**Supplementary Table 2**), as in previous MR studies^13,14^. Each microbial feature had an average of 44 genetic variants (**Fig. 2a** and **Supplementary Table 3**). The corresponding genetic variants explained microbial features to a median value of 24.9%, e.g. 45.5% of the microbial metabolic pathway for succinate consumption and 44.6% of *Phascolarctobacterium succinatutens* (an asaccharolytic, succinate-utilizing bacterium), while only 6.8% of genus *Edwardsiella* (**Supplementary Table 3**). The phenotypic (relative abundance) variance of five genera (*Bilophila, Oscillibacter, Faecalibacterium, Megasphaera* and *Bacteroides*) could be explained over 35% by their corresponding independent genetic variants (**Fig. 2b**). Thus, although human genetic associations (array data) have been reported to explain only 10% or 1.9% of the gut microbiota^7,10^, the suggestive associations from the current M-GWAS study could be highly predictive of certain gut taxa and functions.

**Figure 2.**
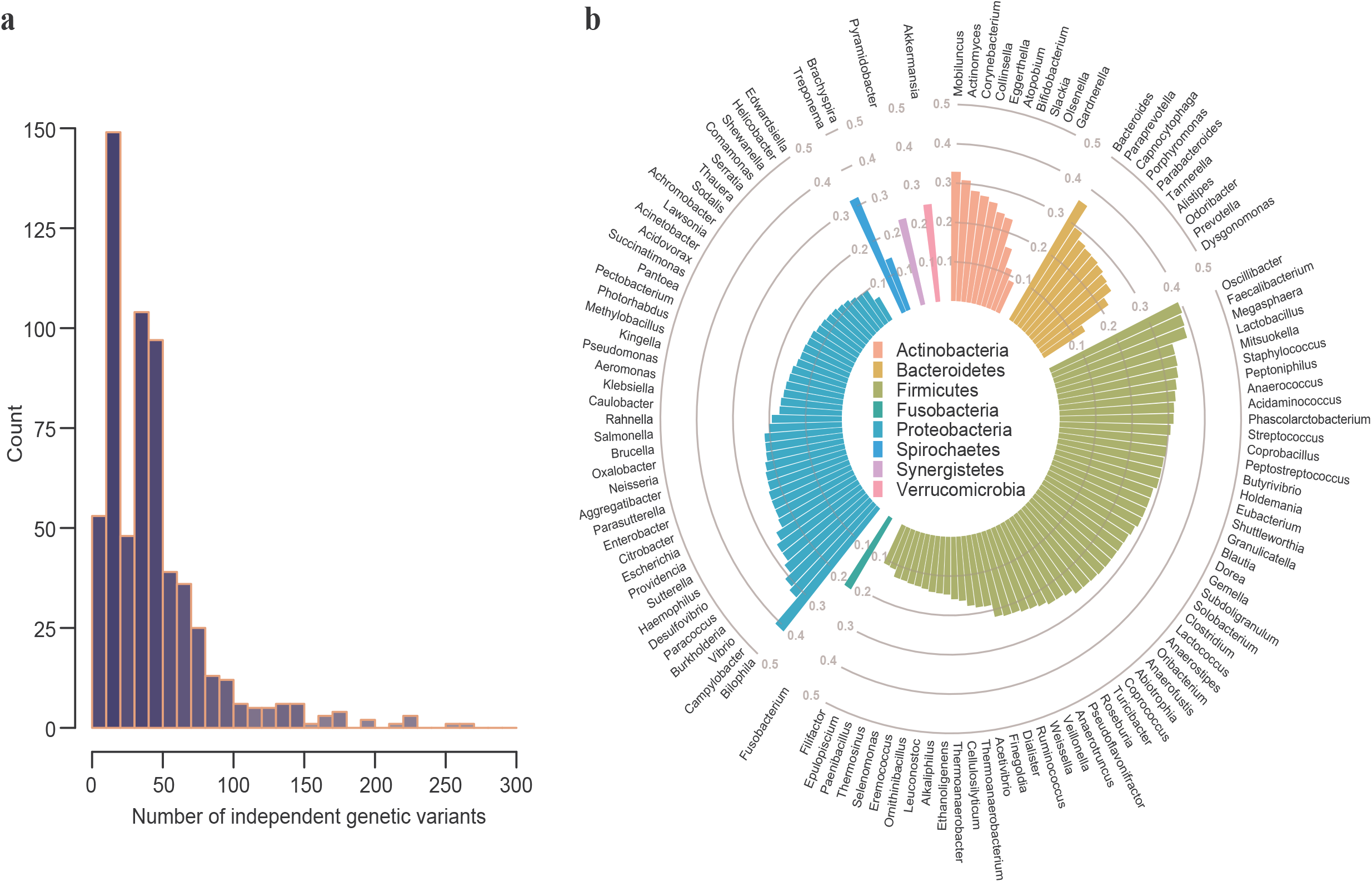
Independent genetic variants and their explained variance of microbial features. **a**, Density plot showing the distribution of number of independent genetic variants that associated with each microbial feature (r^2^ < 0.1 and *P* < 1 × 10^−5^), as calculated by GWAS analysis for a total of 500 unique microbial features. The *x*-axis indicates the number of independent genetic variants for each microbial feature (taxon or GMM). The *y*-axis indicates the number of microbial features under a given number of independent predictors. **b**, Variance explained by the corresponding independent genetic variants for each microbial feature. The polar bar plot indicates how much the independent genetic variants of each common genus (appeared at least 50% of samples) explained for their phenotypic variance (relative abundance of each genus). Genera were classified according to their respective phylum, which are marked with different colors. The *h*^2^ was calculated using REML method in GCTA tools.

For better confidence in these suggestive associations, we sequenced a replication cohort of 1,430 individuals from multiple cities in China (also shotgun metagenomic sequencing for stool samples, but about 8× whole-genome sequencing for human genome; **Supplementary Fig. 1c,d**). Among the 22,293 independent associations identified in the discovery cohort with *P* < 10^−5^, 4,876 variants were not available in the low-depth replication dataset and 87.6% of them were not common variants (MAF < 0.05). We were able to replicate 2,324 of the remaining 17,417 independent associations in the same effect direction of minor allele (*P* < 0.05, **Supplementary Table 2**), indicating that the associations were not random false positives. The fraction of associations replicated in the same direction (*P* < 0.05) using the suggestive cut-off of *P* < 10^−5^ (2,324/17,417) was not lower than the more stringent cut-offs (54/625 of the *P* < 5 × 10^−8^, and 2/28 of the *P* < 10^−10^). Two well-replicated signals from the study-wide threshold were rs1461780285 in the *ABO* blood group associated with module MF0007: lactose and galactose degradation (*P_discovery_* = 2.10 × 10^−12^ and *P_replication_* = 1.09 × 10^−10^; **Supplementary Fig. 3a,b**) and rs142693490 near the *LCORL* gene (implicated in spermatogenesis, body frame and height) associated with MF0034: alanine degradation II (*P_discovery_* = 1.28 × 10^−12^ and *P_replication_* = 0.014; **Supplementary Fig. 3c,d**). Genetic variant rs1461780285 is in strong linkage disequilibrium (LD, *r*^2^ = 0.99) with multiple SNPs (rs507666, rs532436, rs651007, rs579459 and rs579459) in the *ABO* gene. These SNPs located in a block were found to be associated with metabolites levels in both this study and previous studies, especially for serum alkaline phosphatase levels (**Supplementary Table 4**). In addition, we were able to replicate several previously reported microbial signals^6–10^, especially for rs12354611 and *Bacteroides stercoris* (*P* = 8.64 × 10^−6^; **Supplementary Table 5** and **Supplementary Note**).

### Blood metabolic traits associated with human genetic variants

On the other hand, plasma metabolite levels are also known to associate with host genetic variants (**Fig. 1**). We thus performed whole genome-wide association tests for each of the 112 metabolites, with log-transformed relative concentration. We identified a total of 174 associations involving 158 loci that independently associated with one or more of the 112 metabolites at genome-wide significance (*P* < 5 × 10^−8^). With a more conservative Bonferroni-corrected study-wide significant *P* value of 4.5 × 10^−10^ (= 5 × 10^−8^/112 metabolites), we identified 39 associations with metabolites involving 28 genomic loci (**Supplementary Table 6**). These included previously well-established associations such as the *UGT1A* family associated with serum total bilirubin^15,16^ and *ASPG* associated with asparaginate^15^.

According to the suggestive threshold of *P* < 10^−5^, we identified 6,541 metabolic quantitative trait loci (mQTLs), of which 361 were associated with two or more metabolites (**Supplementary Table 6**). The average number of genetic variants was 58 for each metabolic trait (**Fig. 3a** and **Supplementary Table 7**). The percentage of variance explained by the corresponding genetic variants ranged from 13.3% (red blood cell distribution) to as high as 48.3% (blood mercury concentration) and 45.9% (blood alpha-fetoprotein value), with a median value of 28.6% (**Fig. 3b**). Among these, 268 variants or their proxy variants (*r*^2^ > 0.6; distance < 1 Mb) have been reported in the GWAS catalog^17^ (**Supplementary Table 8**). Some variants were associated with diseases in the GWAS catalog such as chronic kidney disease, Alzheimer’s disease, coronary artery disease, Crohn’s disease, ovarian cancer, breast cancer and gastric cancer.

**Figure 3.**
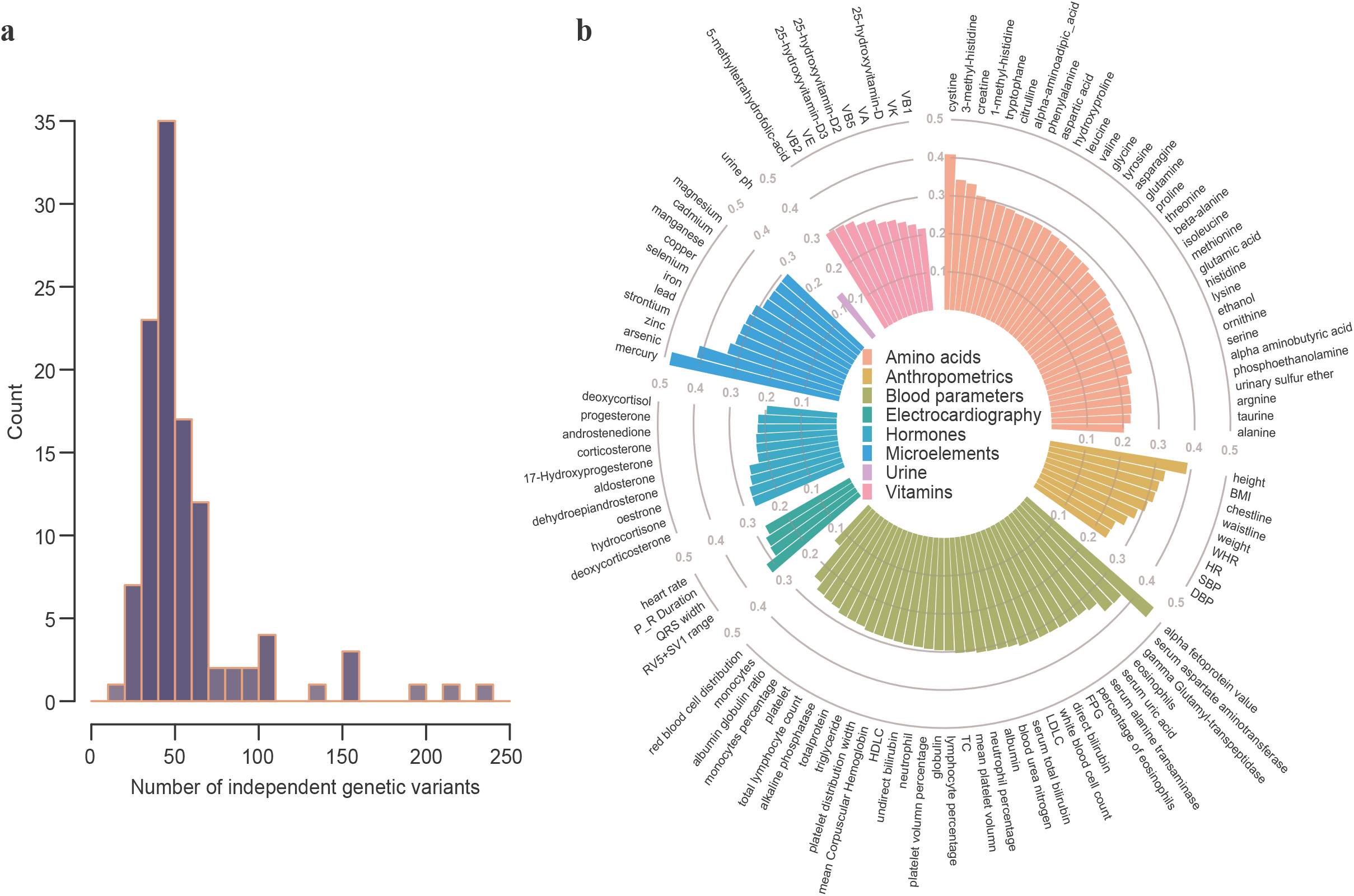
Independent genetic variants and their explained variance of metabolic traits. **a**, Density plot showing the distribution of number of independent genetic variants that associated with each metabolic trait (r^2^ < 0.1 and *P* < 1 × 10^−5^), as calculated by GWAS analysis for a total of 112 metabolic traits. The *x*-axis indicates the number of independent genetic variants for each metabolic trait. The *y*-axis indicates the number of metabolic traits under a given number of independent predictors. **b**, Variance explained by the corresponding independent genetic variants for each metabolic trait. The polar bar plot indicates how much the independent genetic variants of each metabolic trait explained for their phenotypic variance. Each metabolic trait was classified into different catalogs which were marked with different colors. The *h*^2^ was calculated using REML method in GCTA tools.

Among the 6,541 suggestive mQTLs identified in the 4D-SZ discovery cohort with *P* < 10^−5^, 5,088 variants were covered by the replication dataset. 717 and 31 were replicated at nominal (*P* < 0.05) and suggestive significance (*P* < 10^−5^), respectively, in the same effect direction of minor allele (**Supplementary Table 6**). Especially for the 174 genome-wide and 39 study-wide significant associations, we could replicate 51 and 29 associations in the same direction (*P* < 0.05), respectively. The top associations confirmed by the low-depth genomes (*P* < 4.5 × 10^−10^ both in discovery and replication cohorts) included: *FECH* associated with manganese; *UGT1A* family associated with serum total bilirubin as well as direct and indirect (unconjugated) bilirubin; *ASPG* associated with asparagine; *CPS1* associated with glycine; and *APOE* associated with low density lipoprotein (**Supplementary Fig. 4**). Overall, the accurate identification of genetic determinants and the high variance explained for both microbial features and blood metabolites are optimal for MR analysis to investigate causality.

### From observational correlation to Mendelian randomization

As a prerequisite for strong causality, we investigated the correlation between relative abundances of 500 unique fecal microbial features (taxa and functional modules) and 112 host metabolic traits using multivariate linear regression. After adjustment for gender and age, we observed 457 significant associations (false discovery rate (FDR) corrected *P* < 0.05; **Supplementary Table 9** and **Online Methods**). Three metabolites, glutamic acid, 5-methyltetrahydrofolic acid (5-methyl THF, active form of folic acid) and selenium, were associated with the largest number of microbial features (58, 40 and 38, respectively; **Supplementary Fig. 5**). These associations extend findings from various studies and suggest quantitative relationships between gut microbial taxa/functions and plasma metabolites.

As we were fortunate to have all the data in the same cohort, we first performed one-sample MR analyses to identify causal relationships for the 457 observational correlations in the discovery cohort consisting of 1,539 individuals with both metabolic and microbiome traits. We found 58 significant causal effects, of which 17 showed causal effects for gut microbial features on blood metabolic traits and the other 41 showed causal effects for blood metabolic traits on gut microbial features (*P* < 1.09 × 10^−4^ = 0.05/457; **Fig. 4** and **Supplementary Table 10**). Only 4 of these were bidirectional. By applying one-sample MR analyses to the replication dataset of 1,006 low-depth genomes as well as metabolic and microbiome traits from individuals in different cities, we could replicate 43 of the 58 causal relationships (in the same direction and *P* < 0.05; **Supplementary Table 10**), indicating that the effects were not random false positives.

**Figure 4.**
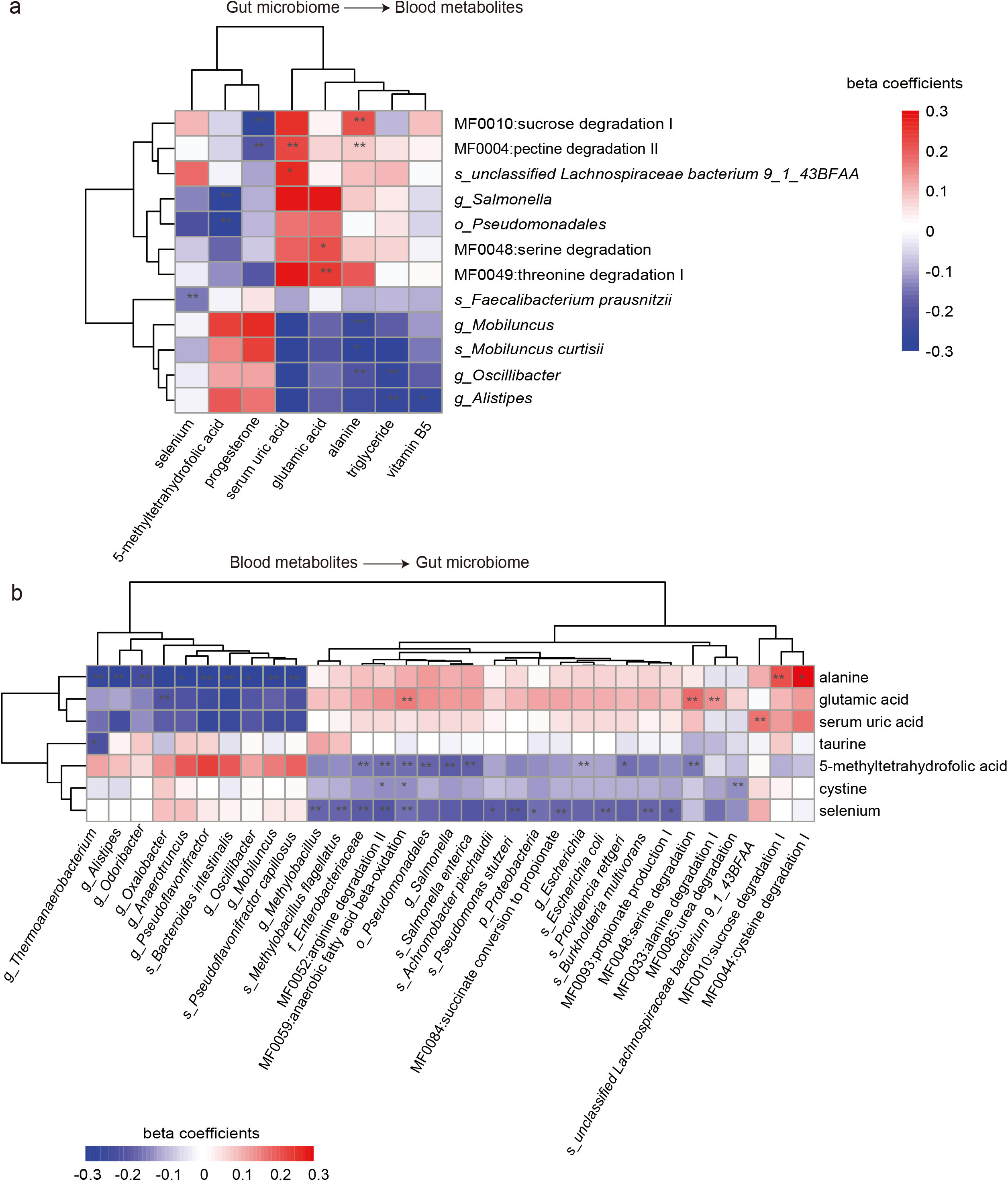
Identifying 58 causal relationships for the microbial features and metabolic traits. **a**, Plot showing the causal effects of 12 specific microbial features on 8 metabolic traits involved in the 17 causal associations from gut microbiome to blood metabolites. **b**, Plot showing the causal effects of 7 blood metabolites on 33 microbial features involved in 41 causal associations from blood metabolites to gut microbiome. Cells marked with “**” represent 43 of the 58 associations identified in discovery cohort that were also replicated in replication cohort, while “*” represent the other 15 only significant in discovery cohort. Cells are colored according to the beta coefficients from one-sample MR analysis, with red and blue corresponding to positive and negative associations, respectively.

Moreover, we also used six different two-sample MR methods, which are more commonly performed when only summary statistics are available from two different cohorts, to analyze our data both in the discovery cohort (summary data for 2,002 samples with metabolic traits and 1,539 samples with microbial features) and the replication cohort (summary data for 1,430 samples with metabolic traits and 1,006 samples with microbial features). The one-sample MR and the two-sample MR analyses showed highly consistent results, and the Spearman’s correlation for beta coefficients between one-sample and two-sample MR reached 0.767 for the discovery cohort (*P* < 2.2 × 10^−16^). The 58 causal associations identified by one-sample MR were also significant in the two-sample MR analyses. An additional 14 causal associations were identified by the two-sample MR analyses (**Supplementary Table 11**), possibly due to the larger cohort size. We also examined the presence of horizontal pleiotropy by using the MR-PRESSO Global test^18^. Only one causal association (the negative effect of selenium on the abundance of *Methylobacillus flagellates, P*_MR-PRESSOGlobaltest_ = 0.01; **Supplementary Table 9**) showed pleiotropy, while all the other 71 causal relationships showed no evidence of pleiotropy (*P* > 0.05). Thus, our MR analyses identified robust causal relationships between blood metabolic traits and specific features of the gut microbiome.

### Effects of the gut microbiome on blood metabolic traits

As some of the MR-identified relationships appeared linked, we performed hierarchical clustering for the 12 microbial features and 8 blood metabolites involved in the 17 causal relationships from the gut microbiome to blood metabolites, which formed two clusters. One cluster involved decreasing the plasma levels of triglyceride and alanine by gut microbial taxa or functional modules, and the other involved decreasing the levels of 5-methyltetrahydrofolic acid or progesterone, but increasing serum uric acid or plasma glutamic acid by gut microbial features (**Fig. 4a**). Reassuringly, the species *Mobiluncus curtisii* was clustered next to its corresponding genus *Mobiluncus*, and modules including serine degradation and threonine degradation, sucrose degradation and pectin degradation, were likewise next to one another.

The most significant causal effect was *Oscillibacter* on decreasing blood triglyceride concentration (**Fig. 5a-c**), and to a lesser extent on lowering body-mass index (BMI) and waist-hip ratio (WHR), whereas the effect with plasma alanine was bidirectional. Using 134 genetic variants to construct a polygenic risk score (PRS) (134 genetic variants and the constructed PRS explained 39.3% and 49.6% of the phenotypic variance, respectively, **Fig. 3b** and **Supplementary Table 10**) for one-sample MR analysis in the discovery cohort, we estimated that each 1-s.d. increase in the abundance of *Oscillibacter* would generate a 0.261 mmol/L decrease in triglyceride concentration (*P* = 2.53 × 10^−10^), a 0.161 kg/m^2^ decrease in BMI (*P* = 1.33 × 10^−4^) and 0.126 ratio decrease in WHR (*P* = 2.73 × 10^−3^). This causal relationship was robust when four two-sample MR tests were performed (*P*_GCTA-GSMR_ = 4.34 × 10^−11^, *P*_Inverse_variance_weighted_ = 2.45 × 10^−15^, *P*_weighted-median_ = 1.22 × 10 and *P*_MR-Egger_ = 1.35 × 10^−5^) (**Fig. 5c**), and there was no evidence of horizontal pleiotropy (*P*_MR-PRESSOGlobaltest_ = 0.18; **Supplementary Table 11**). The reverse MR analysis (testing the effect of genetic predictors of triglyceride on *Oscillibacter* abundance) was significant but did not reach the multiple test corrected significance (10^−4^ < *P* < 0.05). *Oscilibacter* is a Gram-negative Clostridial bacterium, phylogenetically close to *Oscillospira*^19^, which could produce valerate or butyrate. In addition, higher relative abundance of *Alistipes* was also associated with decreased blood triglyceride concentration (*P* = 8.31 × 10^−8^, **Fig. 5d**). At the species level, both *A. shahii (P* = 1.37 × 10^−6^) and *unclassified Alistipes sp. HGB5 (P* = 3.36 × 10^−5^) showed negative effects on blood triglyceride. The effects of both *Oscilibacter* and *Alistipes* for lowering blood triglyceride concentration were confirmed in the replication cohort (*P* = 3.39 × 10^−4^ and *P* = 2.88 × 10^−4^, respectively; **Supplementary Tables 10** and **11**). These findings support the decrease in relative abundances of *Oscillibacter* and *Alistipes* in obese individuals compared to individuals with normal BMI reported in previous studies^20–22^, suggesting that these bacteria are promising supplementation agents for individuals of a suitable genetic background.

**Figure 5.**
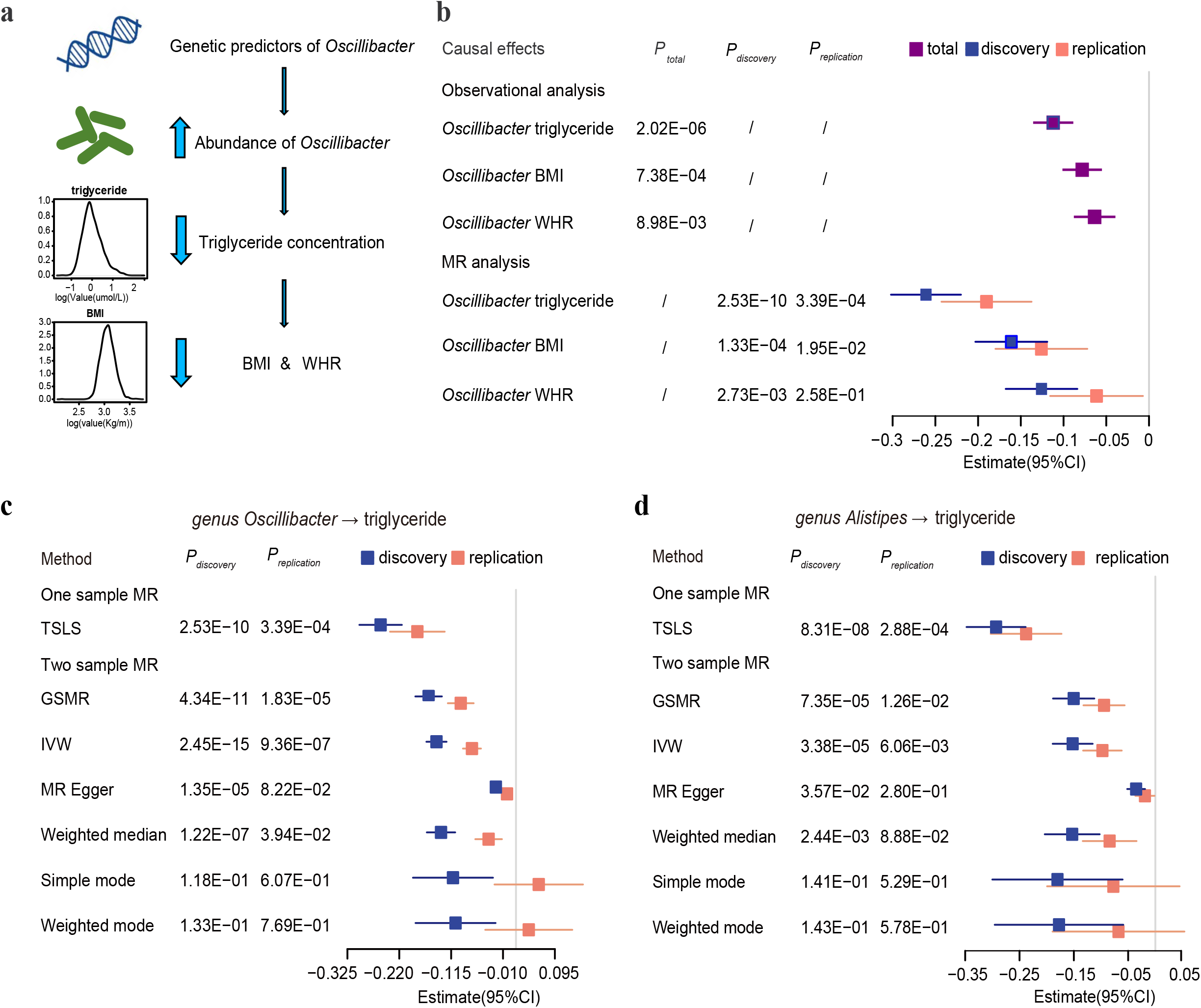
Causal effects of genus *Oscillibacter* and *Alistipes* on decreasing blood triglyceride concentration. **a**, Schematic representation of the MR analysis results: genetic predisposition to higher abundance of *Oscillibacter* is associated with decreased blood triglyceride concentration, and to a lesser extent for lowering body mass index (BMI) and waist-hip ratio (WHR). **b**, Forest plot representing the effect of per 1-s.d. increase in *Oscillibacter* abundance on blood triglyceride, BMI and WHR, as estimated using observational and MR analysis, respectively. Observational correlation analysis was performed using a multivariate linear model in a total of 2,545 samples (purple). One-sample MR analysis was carried out by using a PGS constructed by up to 134 genetic predictors as an instrumental variable, as estimated in discovery cohort (blue) and replication cohort (red), respectively. The beta estimates and 95% confidence interval (CI) values as well as *P* values from both the observational and one-sample MR analysis were listed. **c**,**d**, Forest plots representing the MR estimates and 95% CI values of the causal effects of *Oscillibacter* (**c**) and *Alistipes* (**d**) on triglyceride level, respectively, as estimated using a one-sample MR and six different two-sample MR methods both in discovery cohort (blue) and replication cohort (red). The *P* values calculated by each MR method are also listed.

The gut microbiome potential for pectin degradation II (42.6% of the variance explained by GRS) showed a handful of significant MR hits with blood traits (**Fig. 4a**), including positive effects on alanine (*P* = 8.57 × 10^−5^) and serum uric acid (*P* = 1.34 × 10^−6^), and negative effects on progesterone (*P* = 6.68 × 10^−7^). *Bacteroidetes* and *Fusobacteria* were the only two phyla that positively correlated with the abundance of pectin degradation II (Spearman rank correlation, ρ = 0.48 and 0.15, respectively), which included the two previously reported pectin-degrading species *Bacteroides thetaiotaomicron* and *Fusobacterium varium*^23,24^. In the 4D-SZ cohort, *F. varium* correlated with pectin degradation II (Spearman’s correlation, ρ = 0.12) and increased blood alanine (*P* = 0.02) and serum uric acid (*P* = 0.04); *B. thetaiotaomicron* correlated with pectin degradation II (Spearman’s correlation, ρ = 0.21) but showed no detectable effect on alanine or uric acid (*P* > 0.05; **Supplementary Fig. 7a,b,d**). Instead, *B. dorei*, the bacterial species most strongly correlated with pectin degradation II (Spearman rank correlation, ρ = 0.32, **Supplementary Fig. 7c**), positively contributed to alanine (*P* = 0.05) and serum uric acid levels (*P* = 3.40 × 10^−4^; **Supplementary Fig. 7d**).

### Effects of blood metabolites on gut microbial features

For the 41 causal relationships from blood metabolic traits to gut microbial features (one-sample MR, **Supplementary Table 10**), hierarchical clustering revealed two clusters, one mostly involved decreasing abundance of bacteria by plasma alanine or glutamic acid, and the other involved decreasing abundance of bacteria by selenium or 5-methyltetrahydrofolic acid (**Fig. 4b**). *F. prausnitzii* showed a negative effect on plasma selenium (**Fig. 4a**), while plasma selenium showed negative effects on gut *Proteobacteria* such as Enterobacteriaceae (e.g. *Escherichia coli, P* = 3.79 × 10^−5^), *Pseudomonas stutzeri (P* = 1.06 × 10^−6^), and modules such as arginine degradation II (*P* = 2.65 × 10^−6^), succinate conversion to propionate (*P* = 3.55 × 10^−5^), and anaerobic fatty acid beta oxidation (*P* = 9.71 × 10^−5^) (**Fig. 4b**).

Bacteria from the phylum *Proteobacteria* were negatively affected by not only selenium but also 5-methyltetrahydrofolic acid (**Fig. 4b**). We directly verified the effect of 5-methyltetrahydrofolic acid on *Escherichia in vitro*. Supplementing 5-methyltetrahydrofolic acid in growth media indeed slowed down the growth of a strain of *Escherichia coli* AM17-9 compared to lower concentrations or absence of 5-methyltetrahydrofolic acid (**Supplementary Fig. 8**).

A handful of bacteria were also affected by glutamic acid. The negative influence of glutamic acid (48 variants with suggestive associations and the constructed PRS explained 24.9% and 25.4% of the phenotypic variance, respectively) on the genus *Oxalobacter (P* = 1.56 × 10^−6^) may help explain the lower prevalence of *Oxalobacter* in developed countries, besides the lower intake of oxalate and antibiotic use^25^. Whether limiting glutamic acid could raise *Oxalobacter* and prevent kidney stones remains to be tested. Glutamic acid negatively affected melibiose degradation (to glucose, galactose, *P* = 2.05 × 10^−5^ from two-sample MR), but showed positive effects on alanine degradation I (*P* = 5.46 × 10^−5^), anaerobic fatty acid beta-oxidation (*P* = 9.36 × 10^−5^), and bidirectional positive effect on serine degradation (*P* = 6.85 × 10^−7^ for serine degradation to glutamic acid and *P* = 9.90 × 10^−6^ for glutamic acid to serine degradation, respectively).

### Causal relationships between the gut microbiome and diseases

We further investigated the effects of the 72 significant causal relationships (**Supplementary Table 11**) involving 40 microbial features and 12 metabolic traits on diseases by performing two-sample MR analysis using gut microbiome GWAS summary data in this 4D-SZ cohort, together with blood quantitative traits and diseases GWAS summary statistics from Biobank Japan^26^ (**Fig. 1** and **Supplementary Table 12**), given that Japanese people have a genetic architecture similar to Chinese. Only routine blood parameters but no amino acids, hormones or microelements were included in the Biobank Japan study. Thus, only five of the 72 causal associations, involving triglyceride and serum uric acid were available for further investigation in the Biobank Japan data. The relationship between unclassified Lachnospiraceae bacterium 9_1_43BFAA and uric acid was reciprocal in the 4D-SZ cohort, and we could replicate the causal effect of uric acid on increased unclassified Lachnospiraceae bacterium 9_1_43BFAA abundance in the Japanese cohort, whereas the reciprocal effect, i.e. potential effect of unclassified Lachnospiraceae bacterium 9_1_43BFAA on uric acid was not replicated, possibly due to lack of variants in the genotyped Japanese cohort (15 instead of 67, **Supplementary Table 13**). The other three associations were not replicated, possibly for the same reason. For example, genus *Oscillibacter* had 135 variants with *P* < 10^−5^ in our summary data but only 15 were available in the Biobank Japan summary data.

MR inference using our gut microbiome M-GWAS summary data and disease GWAS summary statistics from Biobank Japan found that *Alistipes*, which showed negative effects on blood triglyceride in the 4D-SZ cohort, lowered the risks of cerebral aneurysm (**Supplementary Table 14**; *P* = 4.61 × 10^−4^) and hepatocellular carcinoma (*P* = 0.045) in the Biobank Japan cohort. According to the genetic associations we identified for *Proteobacteria*, we were able to see in Biobank Japan disease data that *Proteobacteria* increased the risk of T2D (**Fig. 6a**; *P* = 7.61 × 10^−4^, two-sample MR), congestive heart failure (*P* = 0.003) and colorectal cancer (*P* = 0.047). This is consistent with MWAS findings mainly for Enterobacteriaceae^1^ and suggest that the metabolites identified above (5-methyltetrahydrofolic acid, selenium) might help prevent the diseases. Folic acid is indeed recommended for heart diseases^27^. In addition, *Escherichia coli* increased the risk of urolithiasis (**Fig. 6b**; *P* = 0.009) and hepatocellular carcinoma (*P* = 0.04) but decreased interstitial lung disease risk (*P* = 0.007). Similarly, *Salmonella enterica* increased prostate cancer risk but decreased interstitial lung disease risk. The Pseudomonadales order was the only microbial feature showing a positive effect on pulmonary tuberculosis. The denitrifying bacteria *Achromobacter* increased the risk of atopic dermatitis (*P* = 0.005), gastric cancer (*P* = 0.008), esophageal cancer (*P* = 0.027) and biliary tract cancer (*P* = 0.034). *Bacteroides intestinalis*, which was reported to be relatively depleted in patients of atherosclerotic cardiovascular disease^28^, was found here to increase with potassium, and *B. intestinalis* showed a negative effect on epilepsy. *Streptococcus parasanguinis* had a positive effect on colorectal cancer and posterior wall thickness (echocardiography), consistent with MWAS studies^1,28,29^. These results illustrated the potential significance of the gut microbiome-blood metabolite relationships in understanding and preventing cardiometabolic diseases and cancer.

**Figure 6.**
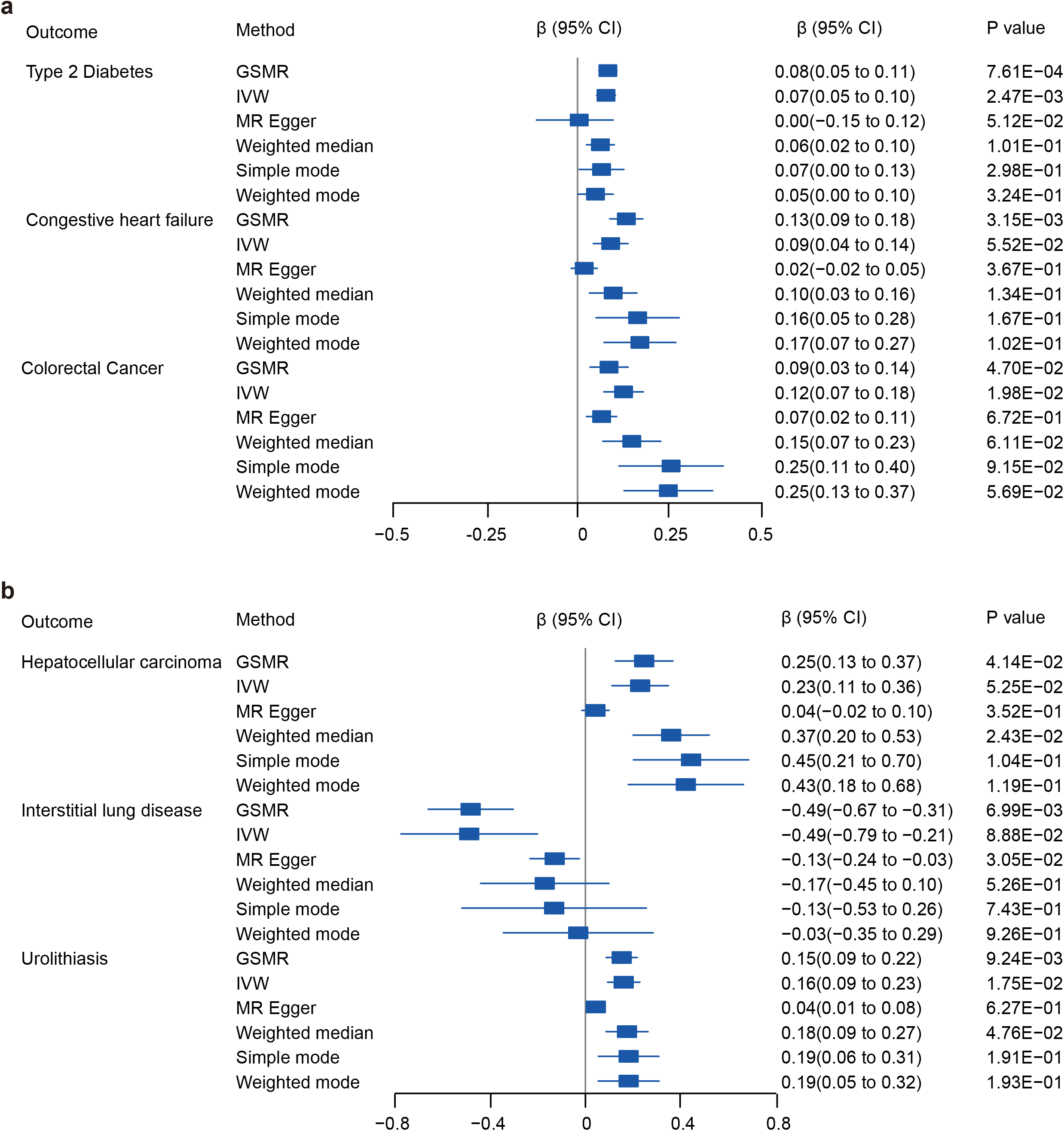
Causal effects of *Proteobacteria* and *Escherichia coli* on diseases. **a**,**b**, Forest plots representing the MR estimates and 95% CI values of the causal effects of *Proteobacteria* (**a**) and *Escherichia coli* (**b**) on diseases, as estimated using six different two-sample MR approaches (GSMR, IVW, MR Egger, Weighted median, Simple mode and Weighted mode). We show the estimated β values (95%CI) and *P* values, as estimated by each MR approach by using the diseases’ summary statistics from Biobank Japan and the gut microbiome GWAS summary data from this discovery cohort with high-depth WGS.

## Discussion

In summary, we identified many genetic loci associated with microbial features and metabolic traits, and found 58 causal relationships between the gut microbiome and blood metabolites using one-sample bidirectional MR. 43 out of the 58 one-sample MR signals could be replicated in a low-depth genome sequenced cohort also from China. Two-sample MR replicated the 58 causal relationships in the same direction and identified an additional 14 causal relationships. Two-sample MR using summary statistics from Biobank Japan identified effects of gut microbial features on diseases, suggesting potential applications of microbiome intervention in cardiometabolic, kidney and lung diseases, and cancer. While mechanistic investigations using germ-free mice and reference bacterial strains have been popular, our data-driven analyses underscore the clinical relevance of gut microbes that have not been extensively cultured and characterized, e.g. *Oscilibacter* and *Alistipes* for lowering triglyceride concentration and a number of disease risks, which may be particularly relevant for East Asian regions undergoing rapid changes in lifestyle and disease profiles.

Although associations between the gut microbiome and blood features such as amino acids and vitamins have been known for some time, our MR analyses could inspire more mechanistic and interventional studies. The unique data available from the 4D-SZ cohort allowed appreciation of overlooked features such as selenium. Interestingly, while selenium showed adverse effects on Gammaproteobacteria, *F. prausnitzii* showed a negative effect on plasma selenium. Nitrogen is a limiting resource for many ecosystems. In the modern human gut microbiome without high intake of nitrite, proteins are probably the major source of nitrogen^30^, and the glutamate-glutamine reservoir is a key buffering mechanism for the inflammatory potential of excess amines^31–34^. The increase in *Proteobacteria* and decrease in Oxalobacteraceae observed in these Chinese individuals no more than 30 years old on average could potentially explain susceptibility to cardiometabolic and kidney diseases later in life. The bidirectional link between strontium and *Streptococcus parasanguinis* implies an interplay between water source and cardiovascular diseases^28,35^.

Metabolism of polysaccharides that cannot be directly digested by the host is an important function of the colonic microbiome. We found degradation of pectin (or sucrose) to negatively affect progesterone level. This is an interesting possibility to provide scientific support for traditional dietary advice for pregnant women to ensure full-term pregnancy. Hyperuricemia and gout is a growing epidemic in East Asia, and soft drinks containing fructose are a strong factor that is no less important than beer and meat^36^. Gut microbial (*Bacteroides*, *Fusobacterium*) pectin degradation module positively contributed to circulating levels of alanine and uric acid. Further studies on the trans-kingdom metabolic flux of carbon and nitrogen would be necessary for personalized management of uric acid and alanine levels.

For the nascent field of M-GWAS and microbiome MR, there are also a lot of opportunities for methodological development by statistical experts. Low-frequency microbes are common in an individual’s gut and could play physiological or pathological roles^1,37^. Our MR results for gut microbial species were supported by MR for higher taxonomic units such as genus or phylum (**Fig. 4** and **Supplementary Table 11**). Yet, the *P* values were sometimes more significant for the larger taxa, suggesting similar functions contributed by other species. Functional redundancy in the microbiome has been discussed ever since the beginning of the microbiome field^38,39^, and here we identified study-wide significant host genetic associations with gut microbial functional modules, and causal effects of other gut microbial functional modules on host levels of circulating metabolites. Distribution of the microbiome taxonomic or functional data constitutes another layer of consideration, in addition to the human allele frequencies. Gathering a more homogenous cohort could enable identification of signals in a relatively small cohort, while corrections for comparing different populations might involve host-microbiome interactions. As the gut microbiome can be influenced by medication^40^, and heritability for most traits is higher in younger individuals^41^, healthy young adults are probably preferable for M-GWAS studies, while microbiome-drug interactions in older individuals could be an important direction for MR studies.

Overall, our data-driven approach underscores the great potential of M-GWAS and MR for a full picture of the microbiome, which can be mechanistically illuminating and are poised to help focus intervention efforts to mitigate inflammation and prevent or alleviate complex diseases.

## Acknowledgements

We sincerely thank the support provided by China National GeneBank. We thank all the volunteers for their time and for self-collecting the fecal samples using our kit. We are very grateful to Professor Karsten Kristiansen (Department of Biology, University of Copenhagen, Denmark; BGI-Qingdao, BGI-Shenzhen, China) for his support for the joint PhD program.

## Author Contributions

H.J. and T.Z. conceived and organized this study. J.W. initiated the overall health project. X.X., H.Y., S.Z., Y.H., W.L. and Y. Zong contributed to organization of the cohort sample collection and questionnaire collection. H.L. led the DNA extraction and sequencing. X.Q., J.Z., and R.W. generated the metabolic data. X. Liu, T.Z., and X.T. processed the whole genome data. Y. Zou, X. Lin, Z.Z., H.Z., L.T., Q.W., Z.J. and L.X. processed the metagenome data. X. Liu and X.T. performed the Mendelian randomization analyses. X. Liu and H.J. wrote the manuscript. All authors contributed to data and text in this manuscript.

## Methods

### Study participants

All adult Chinese individuals included in this study were recruited for a multi-omic study, with some volunteers having samples from as early as 2015, which would constitute the time dimension in ‘4D’. The discovery cohort was recruited during a physical examination from March to May in 2017 in the city of Shenzhen, including 2,002 individuals with blood samples, of which 1,539 had fecal samples. All these individuals were enlisted for high-depth whole genome and whole metagenomic sequencing. For replication, blood samples were collected from 1,430 individuals, of which 1,006 had fecal samples. The replication cohort was designed in the same manner but organized at smaller scales in multiple cities (Wuhan, Qingdao, etc.) in China. The protocols for blood and stool collection, as well as the whole genome and metagenomic sequencing, were similar to our previous studies^5,42–44^. For blood samples, buffy coat was isolated and DNA was extracted using HiPure Blood DNA Mini Kit (Magen, Cat. no. D3111) according to the manufacturer’s protocol. Feces were collected with MGIEasy kit, and stool DNA was extracted in accordance with the MetaHIT protocol as described previously^5,42^. The DNA concentrations from blood and stool samples were estimated by Qubit (Invitrogen). 200 ng of input DNA from blood and stool samples were used for library preparation and then processed for paired-end 100-bp and single-end 100-bp sequencing, respectively, using BGISEQ-500 platform^45^.

The study was approved by the Institutional Review Boards (IRB) at BGI-Shenzhen, and all participants provided written informed consent at enrolment.

### High-depth whole genome sequence for discovery cohort

2,002 individuals in discovery cohort were sequenced to a mean of 42× for whole genome. The reads were aligned to the latest reference human genome GRCh38/hg38 with BWA^46^ (version 0.7.15) with default parameters. The reads consisting of base quality <5 or containing adaptor sequences were filtered out. The alignments were indexed in the BAM format using Samtools^47^ (version 0.1.18) and PCR duplicates were marked for downstream filtering using Picardtools (version 1.62). The Genome Analysis Toolkit’s (GATK^48^, version 3.8) BaseRecalibrator created recalibration tables to screen known SNPs and INDELs in the BAM files from dbSNP (version 150). GATKlite (v2.2.15) was used for subsequent base quality recalibration and removal of read pairs with improperly aligned segments as determined by Stampy. GATK’s HaplotypeCaller were used for variant discovery. GVCFs containing SNVs and INDELs from GATK HaplotypeCaller were combined (CombineGVCFs), genotyped (GenotypeGVCFs), variant score recalibrated (VariantRecalibrator) and filtered (ApplyRecalibration). During the GATK VariantRecalibrator process, we took our variants as inputs and used four standard SNP sets to train the model: (i) HapMap3.3 SNPs; (ii) dbSNP build 150 SNPs; (iii) 1000 Genomes Project SNPs from Omni 2.5 chip; and (iv) 1000G phase1 high confidence SNPs. The sensitivity threshold of 99.9% to SNPs and 99% to INDELs were applied for variant selection after optimizing for Transition to Transversion (TiTv) ratios using the GATK ApplyRecalibration command. After applying the recalibration, there were 60,978,451 raw variants left, including 55 million SNPs, and 6 million INDELs.

We applied a conservative inclusion threshold for variants: (i) mean depth >8×; (ii) Hardy-Weinberg equilibrium (HWE) *P* > 10^−5^; and (iii) genotype calling rate > 98%. We demanded samples to meet these criteria: (i) mean sequencing depth > 20×; (ii) variant calling rate > 98%; (iii) no population stratification by performing principal components analysis (PCA) analysis implemented in PLINK^49^ (version 1.9); and (iv) excluding related individuals by calculating pairwise identity by descent (IBD, Pi-hat threshold of 0.1875) in PLINK. Only 10 samples were removed in quality control filtering. After variant and sample quality control, 1,992 individuals with 6.12 million common (MAF ≥ 5%) and 3.90 million low-frequency (0.5% ≤ MAF < 5%) variants from discovery cohort were left for subsequent analyses.

### Low-depth whole genome sequence for replication cohort

1,430 individuals in the replication cohort were sequenced to a mean of 8× for whole genome. We used BWA to align the whole genome reads to GRCh38/hg38 and used GATK to perform variant calling by applying the same pipelines as for the high-depth WGS data. After completing the joint calling process with CombineGVCFs and GenotypeGVCFs options, we obtained 43,402,368 raw variants. A more stringent process in the GATK VariantRecalibrator stage compared with the high-depth WGS was then used, and the sensitivity threshold of 98.0% to both SNPs and INDELs was applied for variant selection after optimizing for Transition to Transversion (TiTv) ratios using the GATK ApplyRecalibration command. Further, we kept variants with < 10% missing genotype frequency and minor allele count > 5. All these high-quality variants were then imputed using BEAGLE 5^50^ with the 1,992 high-depth WGS dataset as reference panel. We retained only variants with imputation info. > 0.7 and obtained 10,905,418 imputed variants. We further filtered this dataset to keep variants with Hardy-Weinberg equilibrium *P* > 10^−5^ and genotype calling rate > 90%. Similar to what we did for the discovery cohort, samples were required to have mean sequencing depth > 6×, variant call rate > 98%, no population stratification and no kinship. Finally, 1,430 individuals with 5,884,439 high-quality common and low-frequency variants (MAF ≥ 0.5%) from the replication cohort were left for subsequent analysis. To assess the data quality, we sequenced 27 samples with both high-depth and low-depth WGS data and then compared the 5,318,809 variants between them for each individual. The average genotype concordance was 98.66% (**Supplementary Table 15**).

### Metagenomic sequencing and profiling

High-quality whole metagenomic sequencing was performed for 1,539 samples from the discovery cohort and 1,004 samples from the replication cohort with fecal samples available. The reads were aligned to hg38 using SOAP2^51^ (version 2.22; identity ≥ 0.9) to remove human reads. The gene profiles were generated by aligning high-quality sequencing reads to the integrated gene catalog (IGC) by using SOAP2 (identity ≥ 0.95) as previously described^37^. The relative abundance profiles of phylum, order, family, class, genera and species were determined from the gene abundances. To eliminate the influence of sequencing depth in comparative analyses, we downsized the unique IGC mapped reads to 20 million for each sample. The relative abundance profiles of gene, phylum, order, family, class, genus and species were determined accordingly using the downsized mapped reads per sample.

GMMs (gut metabolic modules) reflect bacterial and archaeal metabolism specific to the human gut, with a focus on anaerobic fermentation processes^52^. The current set of GMMs was built through an extensive review of the literature and metabolic databases, inclusive of MetaCyc^53^ and KEGG, followed by expert curation and delineation of modules and alternative pathways. Finally, we identified 620 common microbial taxa and GMMs present in 50% or more of the samples.

### Metabolic trait profiling

Measurements of metabolic traits (anthropometric characteristics and blood metabolites) were performed for all 3,432 individuals during the physical examination in this study. The clinical tests, including blood tests and urinalysis, were performed in licensed physical examination organization. The anthropometric measurements such as height, weight, waistline and hipline were measured by nurses. Age and gender were self-reported. The metabolites, i.e. vitamins, hormones, amino acids and trace elements including heavy metals, were chosen from a health management perspective. Measurements of blood metabolites were performed as described in detail by Jie et al.^39^; blood amino acids were measured by ultra high pressure liquid chromatography (UHPLC) coupled to an AB Sciex Qtrap 5500 mass spectrometry (AB Sciex, US) with the electrospray ionization (ESI) source in positive ion mode using 40 μl plasma; blood hormones were measured by UHPLC coupled to an AB Sciex Qtrap 5500 mass spectrometry (AB Sciex, US) with the atmospheric pressure chemical ionization (APCI) source in positive ion mode using 250 μl plasma; blood trace elements were measured by an Agilent 7700x ICP-MS (Agilent Technologies, Tokyo, Japan) equipped with an octupole reaction system (ORS) collision/reaction cell technology to minimize spectral interferences using 200 μl whole blood; water-soluble vitamins were measured by UPLC coupled to a Waters Xevo TQ-S Triple Quad mass spectrometry (Waters, USA) with the electrospray ionization (ESI) source in positive ion mode using 200 μl plasma; fat-soluble vitamins were measured by UPLC coupled to an AB Sciex Qtrap 4500 mass spectrometry (AB Sciex, USA) with the atmospheric pressure chemical ionization (APCI) source in positive ion mode using 250 μl plasma.

### Observational correlation analyses

As many microbial features (taxonomies and pathways) are highly correlated, we first performed a number of Spearman correlation tests and kept only one member of pairs of bacteria or GMMs showing >0.99 correlation coefficient. This filtering resulted in a final set of 500 unique features (99 GMMs and 401 gut taxa) that were used for analyses. We correlated these 500 microbial features with 112 measured metabolic traits, including 9 anthropometric measurements (BMI, WHR, etc.) and 103 blood metabolites (amino acids, vitamins, microelements, etc.) in the 3,432 individuals. All metabolic traits and microbial features were transformed using natural logarithmic function to reduce skewness of distributions. For each phenotype, we excluded outlier individuals with more than four standard deviations away from the mean. The metabolite measures were then centered and scaled to mean of 0 and standard deviation of 1.

The relationship between metabolic traits and microbial features were evaluated by multivariable linear regression analysis while adjusted for age and gender. After achieving the raw *P* value, we used the *p.adjust()* function in R (v3.2.5)) to perform the multiple test correction and calculated adjusted *P* values with the Benjamini–Hochberg procedure. The results were considered significant when FDR adjusted *P* value was < 0.05. The correlated microbial features and metabolic traits, raw *P* and FDR adjusted *P* values, are included in **Supplementary Table 9**.

### Clustering of microbiome-metabolite associations

To assess the association clusters of 58 identified causal relationships involving the effects of 12 microbial features on 8 metabolic traits and the effects of 7 metabolic traits on 33 microbial features, we performed a hierarchical clustering analysis. Beta coefficients of associations between the microbial features and metabolic traits from one-sample MR analysis were used to construct distance matrices. Complete-linkage hierarchical clustering was used to cluster the metabolites and microbiome traits from the distance matrices using the ‘hclust’ function in R, and the results were visualized as a heatmap.

### Genome-wide association analysis for microbial features

We tested the associations between host genetic variants and gut bacteria using linear or logistic model based on the abundance of gut bacteria. The abundance of bacteria with occurrence rate over 95% in the cohort was transformed by the natural logarithm and the outlier individual who was located away from the mean by more than four standard deviations was removed, so that the abundance of bacteria could be treated as a quantitative trait. Otherwise, we dichotomized bacteria into presence/absence patterns to prevent zero inflation; the abundance of bacteria could then be treated as a dichotomous trait. Next, for 10 million common and low-frequency variants (0.5% ≤ MAF < 5%) identified in the discovery cohort and 5.9 million common and low-frequency variants identified in replication cohort, we performed a standard single variant (SNP/INDEL)-based M-GWAS analysis via PLINK using a linear model for quantitative trait or a logistic model for dichotomous trait. Given the effects of diet and lifestyles on microbial features, we included age, gender, BMI, defecation frequency, stool form, 12 diet and lifestyle factors, as well as the top four principal components (PCs) as covariates for M-GWAS analysis in both the discovery and the replication cohort.

### Genome-wide association analysis for metabolic traits

For each of the 112 anthropometric and metabolic traits, the log_10_-transformed of the median-normalized values was used as a quantitative trait. Samples with missing values and values beyond 4 s.d. from the mean were excluded from association analysis. Each of the 10 million common and low-frequency variants identified in the discovery cohort and the 5.9 million common and low-frequency variants identified in the replication cohort was tested independently using a linear model for quantitative trait implemented in PLINK. Age, gender and the top four PCs were included as covariates.

### Independent predictor and explained phenotypic variance

For each whole genome-wide association result of microbial features and metabolic traits, we first selected genetic variants that showed association at *P* < 1 × 10^−5^ and then performed the linkage disequilibrium (LD) estimation with a threshold of LD *r*^2^ < 0.1 for clumping analysis to get independent genetic predictors. The *P*-value threshold of 1 × 10^−5^ was used for selection of genetic predictors associated with microbial features by maximizing the strength of genetic instruments and the amount of the average genetic variance explained by the genetic predictors in an independent sample. For each microbial feature, we got genetic instruments in discovery cohort using different *P* thresholds, including 5 × 10^−8^, 1 × 10^−7^, 1 × 10^−6^ and 1 × 10^−5^. We tested the strength of these instruments under different *P* thresholds by checking whether they predicted corresponding microbial features in an independent sample (**Supplementary Note**, **Supplementary Table 16** and **Supplementary Fig. 6**), and we observed that the mean value of instrumental F statistics is 3.57 and on average only 0.28% phenotype variance could be explained by instruments on microbial features when using 5 × 10^−8^ as instrumental cut-off. Therefore, we used a more liberal threshold of *P* < 1 × 10^−5^ to select the instruments for microbial features, and the instrumental mean F statistics reached 51.4 (greater than 10), which indicates a strong instrument^54^. The average phenotypic variance explained by instruments on microbial features was 22.6% for the discovery cohort and 5.09% for the replication cohort (**Supplementary Fig. 2**). For consistency, we used the same threshold and procedure for selecting genetic predictors of metabolic traits in both the discovery and the replication cohort. The LD estimation between variants was calculated in 2,002 samples for the discovery cohort and in 1,430 samples for the replication cohort, respectively. For each phenotype, the variance explained by the corresponding independent genetic predictors was estimated using a restricted the maximum likelihood (REML) model as implemented in the GCTA software^55^. We adjusted for age, gender and the top four PCs in the REML analysis.

### One-sample MR analysis

To investigate the causal effects between microbial features and metabolic traits available from the same cohort, we first performed one-sample bidirectional MR analysis in discovery cohort, which included 1,539 individuals with both metabolite and microbiome traits. We specified a threshold of *P* < 1 × 10^−5^ to select SNP instruments and LD *r*^2^ < 0.1 threshold for clumping analysis to get independent genetic variants for MR analysis. Then, an unweighted polygenic risk score (PRS) was calculated for each individual using independent genetic variants from GWAS data. Each SNP was recoded as 0, 1 and 2, depending on the number of trait-specific risk increasing alleles carried by an individual. We performed Instrumental variable (IV) analyses employing two-stage least square regression (TSLS) method^56^. In the first stage, for each exposure trait, association between the GRS and observational phenotype value was assessed using linear regression and predicted fitted values based on the instrument were obtained. In the second stage, linear regression was performed with outcome trait and genetically predicted exposure level from the first stage. In both stages, analyses were adjusted for age, gender and top four principal components of population structure. For each trait, TSLS was performed using ‘ivreg’ command from the AER package in R. We attempted to replicate the causal effects between traits in replication cohort with 1,004 individuals.

### Two-sample MR analysis

To maximize the sample size in MR analysis and confirm the causal effect between microbial features and metabolic traits, we also performed two-sample bidirectional MR analysis using six different methods, including genome-wide complex trait analysis-generalized summary Mendelian randomization (GCTA-GSMR) approach^57^ and five other methods (Inverse-variance weighting (IVW)^58,59^, MR–Egger regression^60,61^, Weighted median^62^, Mode-based estimate (MBE)^63^ including Simple mode and Weighted mode) implemented in “TwoSampleMR” R package as a robust validation. A consistent effect across the six methods is less likely to be a false positive. If the genetic variants have horizontally pleiotropic effects but are independent of the effects of the genetic variants on the exposure, this is known as balanced pleiotropy. If all the pleiotropic effects are biasing the estimate in the same direction (directional pleiotropy), this will bias the results (with the exception of the MR-Egger method). We used the MR-PRESSO (mendelian randomization pleiotropy residual sum and outlier) Global test to estimate for the presence of directional pleiotropy.

We first performed GWAS analysis for every trait and used summary statistics data for MR analysis. Genetic variants with *P* < 1 × 10^−5^ and LD *r*^2^ <0.1 were selected as instrumental variables.

### In vitro growth experiments of *Escherichia coli*

To directly test the interactions between *Escherichia coli* and 5-methyltetrahydrofolic acid, the anaerobic growth of a strain *Escherichia coli* AM17-9 was characterized at different concentrations of 5-methyltetrahydrofolic acid. The *Escherichia coli* AM17-9, isolated from feces of a male, was routinely grown in Luria-Bertani (LB) broth while supplementing 5-methyltetrahydrofolic acid with concentrations of 0, 1 and 2 ng/ml, respectively. The normal concentration of 5-methyltetrahydrofolic acid in human blood ranged from 4.4 ng/ml to 32.8 ng/ml. The growth of *Escherichia coli* AM17-9 was inhibited when supplementing 5-methyltetrahydrofolic acid from 0 to 2 ng/ml. The optical density at 600 nm (OD600) was measured at intervals of two hours using a microplate reader.

### MR analyses for diseases in Biobank Japan

We downloaded summary statistics data for 42 diseases and 59 blood quantitative traits in 212,453 Japanese individuals^26^ (http://jenger.riken.jp/en/result; **Supplementary Table 13**). By combining these data and the gut microbiome GWAS summary data from discovery cohort with high-depth WGS, we performed the two-sample bidirectional MR analysis to investigate the causal effect between the exposure (40 microbial features and 12 metabolic traits that were involved in the 72 significant causal relationships; **Fig. 4**) and the outcome (42 diseases from BioBank Japan) by applying the GSMR method and the other five MR tests as described in the previous paragraph. For consistency, genetic variants with *P* < 1 × 10^−5^ and LD *r*^2^ <0.1 were also selected as instrumental variables for phenotypes in the Biobank Japan study.

## Data availability

The human reference hg38 datasets were publicly available from http://hgdownload.soe.ucsc.edu/goldenPath/hg38/bigZips/. All summary statistics that support the findings of this study including the associations between host genetics and microbiomes, host genetics and metabolites are publicly available from https://ftp.cngb.org/pub/CNSA/data2/CNP0000794/. The release of the data was approved by the Ministry of Science and Technology of China (Project ID: 2020BAT1137). Individual-level data including host genetics, metagenomics and metabolites have been uploaded to the GSA database (https://ngdc.cncb.ac.cn/bioproject/browse/PRJCA005334). Access to individual-level data has to be approved by corresponding authors, and is subject to the policies and approvals from the Human Genetic Resource Administration, Ministry of Science and Technology of the People’s Republic of China. The summary statistics data for 42 diseases and 59 blood quantitative traits in 212,453 Japanese individuals are available from Biobank Japan (http://jenger.riken.jp/en/result).

## Code availability

The host genome reads were aligned to the latest reference human genome GRCh38/hg38 with BWA (v.0.7.15; http://bio-bwa.sourceforge.net/). The alignments were indexed in the BAM format using Samtools (v.0.1.18; http://samtools.sourceforge.net/) and PCR duplicates were marked for downstream filtering using Picardtools (v.1.62; http://broadinstitute.github.io/picard/). The genome variants calling were performed using GATK (v3.8; https://gatk.broadinstitute.org/hc/en-us). The low-depth data were imputed using BEAGLE (v.5.0; https://faculty.washington.edu/browning/beagle/b5_0.html). The metagenome reads were aligned to hg38 using SOAP (v.2.22; http://soap.genomics.org.cn). Quality control, association analyses and genetic risk score analyses were performed in PLINK (v1.90; http://zzz.bwh.harvard.edu/plink/). Variance explained analysis was performed using REML model in GCTA (v.1.26.0; https://cnsgenomics.com/software/gcta). One-sample MR analyses were performed using TSLS method in AER package (v1.2-9; https://www.rdocumentation.org/packages/AER/versions/1.2-9/topics/ivreg). Two-sample MR analyses were performed in GSMR (v1.0.7; http://cnsgenomics.com/software/gsmr/) and TwoSampleMR (v.0.5.6; https://mrcieu.github.io/TwoSampleMR/). All statistics analyses and visualizations were performed in R (v.3.2.5; https://www.r-project.org).

